# Energy trade-offs under fluctuating light govern bioenergetics and growth in *Chlamydomonas reinhardtii*

**DOI:** 10.1101/2025.05.19.654497

**Authors:** Ana Pfleger, Erwann Arc, Shuang Zhang, Palak Chaturvedi, Emma Antoine, Erich Gnaiger, Arindam Ghatak, Leila Afjehi-Sadat, Wolfram Weckwerth, Ilse Kranner, Thomas Roach

## Abstract

Rapidly changing light intensity is a natural challenge that photosynthetic organisms can tolerate. Regulatory mechanisms of light harvesting and alternative electron pathways are critical in dissipating and distributing energy under fluctuating light intensities (FL), but less is known about downstream metabolic regulations. Here, we compared the cellular responses of *Chlamydomonas reinhardtii* grown under FL to cells acclimated to constant high (HL) or low light (LL), either under high (2 %) or low (0.04 %) CO_2_. Under low CO_2_, the physiology of FL cells resembled HL cells and proteomics revealed an induction of the ATP consuming carbon-concentrating mechanism, and photorespiration particularly under FL. High CO_2_ promoted growth under FL, albeit by a lesser extent than under HL and led to higher ATP contents than under low CO_2_. To fuel ATP production under low CO_2_, cells upregulated mitochondrial respiration under FL, while enhanced cyclic electron flow and redox shuttling between intracellular compartments was most evident under FL and LL. Chloroplastic carbon metabolism rapidly responded to light changes, independent of CO_2_ availability, whereas metabolites associated with mitochondrial bioenergetics responded slower, and remained abundant under high CO_2_. The accumulation of enzymes involved in starch synthesis and breakdown under FL, together with the transient accumulation of hexoses and hexose phosphates, indicated that cells relied on sugars as a transient carbon pool to meet changing metabolic demands under FL. We conclude that the interplay between light intensity and CO₂ availability drives critical energy trade-offs, balancing photoprotection, repair, and carbon allocation, that regulate growth under FL.

## 1. Introduction

Algae encounter fluctuating light intensities (FL) in nature and at different time scales, e.g. <1 s from varying cloud cover and sun flecks up to daily (diurnal) and seasonal changes, and also during cultivation due to cell cycling in bioreactors and self-shading (Graham et al., 2017). Changes in light intensity immediately impacts photosystem (PS) activity, thus photosynthetic electron transfer and the trans-thylakoid pH difference (ΔpH), which precedes activation of various regulatory mechanisms (Niyogi, 2000). Photo-regulatory mechanisms include ΔpH-regulated non-photochemical quenching (NPQ), which is induced within seconds to minutes, and protect the PS under high light (HL) & FL conditions. However, the delayed regulation of energy dissipation in response to rapid changes in light intensity, e.g. after a cloud passes in front of the sun, can also lower photosynthetic efficiency (Kromdijk et al., 2016). In *Chlamydomonas reinhardtii*, light-harvesting complex stress related (LHCSR) proteins LHCSR1/3 are the main mediators of high-energy state quenching (qE)-type NPQ (Peers et al., 2009). Other regulatory mechanisms of electron transfer include the energetically wasteful O_2_ photoreduction by photosystem I (PSI), either through the Mehler reaction, or via flavodiiron proteins (FLV), which function in so-called pseudo-cyclic electron flow (PCEF). In cyclic electron flow (CEF), electrons released by PSI re-reduce the photosynthetic electron transfer system (PETS). Two CEF pathways are known in *C. reinhardtii*; one around the PSI and cytochrome *b*_6_*f* complex (Cyt *b*_6_*f*), and the other that uses NAD(P)H dehydrogenase 2 (NDA2). In both routes of CEF, stromal reductants reduce the plastoquinone pool, translocate protons to the lumen and promote ATP synthesis. Collectively, by regulating excitation of the PS (i.e. NPQ) or affecting electron flow, these mechanisms prevent damage to PS reaction centers (i.e. photoinhibition) while balancing the ATP:NADPH ratio for metabolic processes (Peltier et al., 2010). For example, the Calvin Benson Basham (CBB) cycle requires an ATP:NADPH ratio of 3:2, which cannot be achieved by linear electron flow (LEF) alone. Chloroplast to mitochondria electron flow (CMEF), also referred to as redox shuttling, is another important factor for efficient growth and environmental acclimation. CMEF involves the exchange of reducing power through dicarboxylate translocators (OMT) combined with malate dehydrogenases (MDH), often called ‘malate valves’, as well as the export of triose phosphates, which are then consumed by the glycolytic pathway in the cytosol and the TCA cycle in the mitochondria to generate reducing power. This transfer of reducing power via CMEF enables ATP production through oxidative phosphorylation (Shimakawa et al., 2024). Alternative electron flows, especially CEF and PCEF, have been shown to be critical to tolerate FL in *C. reinhardtii* (Jokel et al., 2018), but the importance of CMEF remains unclear (Burlacot, 2023).

Microalgal metabolism is regulated by CO_2_ availability and mechanisms that govern carbon fixation and allocation. Low CO₂ solubility in aquatic environments reduces its availability for photosynthesis (Maberly and Gontero, 2017). In response, algae can activate the so-called CO_2_-concentrating mechanism (CCM), involving ATP-dependent inorganic carbon (C_i_) transporters across all cellular compartments, carbonic anhydrases to enhance C_i_ fixation, and the formation of a starch sheath that surrounds the pyrenoid where Rubisco is located (Mackinder et al., 2017; He et al., 2023). Under CO_2_ limitation, PCEF, CEF and CMEF all may contribute to the additional ATP production required to drive the CCM (Burlacot and Peltier, 2023; Peltier et al., 2024). The CCM helps prevent the oxygenase reaction of Rubisco and production of toxic 2-phosphoglycolate (2PG) that is recycled via photorespiration (Wang et al., 2015). Recycling 2PG through the photorespiratory pathway leads to a loss of 25% of fixed carbon, while consuming 3.5 ATP and 2 NAD(P)H (Fu and Walker, 2023), which further increases the ATP demand beyond what LEF delivers. Hence, the regulation of the ATP/NADPH balance is disrupted under carbon deficiency, an effect that may be further exacerbated under fluctuating light (Fu and Walker, 2023).

The majority of available energy is invested in the synthesis of photosynthetic end products (e.g. reducing sugars, amino acids), accumulation of reserves (e.g. starch, lipids), growth, maintenance and cell division. The partitioning of C is dependent on the cellular energy balance, rate of C assimilation and nutrient availability (Johnson and Alric, 2013; Burlacot et al., 2019). Starch is central to the carbon budget of photosynthetic organisms, acting both as a sink and source. The importance of coordinating starch metabolism with growth is reflected by multiple regulatory interventions in starch biosynthesis and degradation. Post translational regulations of many enzymes involved in starch metabolism enables a rapid induction (seconds to minutes) of starch degradation or synthesis depending on the cell energetic balance (MacNeill et al., 2017). Furthermore, photoautotrophic organisms invest a large fraction of their energy into photosynthetic complexes and rubisco synthesis to be able to achieve photosynthesis. In plants, allocation of C to proteins is light dependent and accounts for up to 2.5 % of net photosynthesis (Tcherkez et al., 2020). In *C. reinhardtii*, cellular protein concentration is a regulator of the cell cycle (Donnan and John, 1983). However, little is known about how CO_2_ and light availability influences protein synthesis in algae.

Switching from a shaded area to direct light exposition may result in an increase in light intensity by over 10-fold (Wang, 2024). So far, research has focused on understanding the response of low light (LL)-acclimated cells to HL, or how cells acclimate to constant LL or HL (Bonente et al., 2011) or in response to high and low CO_2_ availability (Polukhina et al., 2016; Zuliani et al., 2024). Furthermore, intermittent light (i.e. with darkness) imposes a penalty on growth in *C. reinhardtii* that was attributed to de-activation of key enzymes in the CBB cycle (Graham et al., 2017), making the HL phase of FL more stressful (Roach, 2020) However, it is not clear how algae metabolically respond to FL, especially in situations under which light or C_i_ availability may result in photosynthesis limitations. Since algae have a central position in green economy, an in-depth understanding of the response of the core metabolic pathways to these conditions is needed. Accelerating the photosynthetic and metabolic response to light fluctuations may enable new opportunities for algal cultivation, for example in maximizing the production of essential lipids, high value carotenoids and other nutraceuticals. Here, we took a systems-level approach to reveal how contrasting CO_2_ availability influences the interplay between primary metabolism and cellular bioenergetics under FL in governing carbon allocation to growth in *C. reinhardtii*.

## Results & Discussion

### CO2 availability leads to differential physiological acclimation of FL cells to HL

The common wild-type lab strains of *C. reinhardtii* are shade adapted, typical of this soil-dwelling organism (Harris, 2009). For determining the light intensity for photoautotrophic growth conditions, we measured how oxygen fluxes were influenced by light intensity. Gross rate of light-induced O_2_ production saturated at a light intensity of about 250 µmol m^-2^ s^-1^ in cultures acclimated to LL (50 µmol m^-2^ s^-1^) or HL (500 µmol m^-2^ s^-1^) under ambient CO_2_ (Amb; 0.04 %) levels (Supplementary Fig. S1). Whereas similar results were found after LL acclimation under 2 % CO_2_ (LL-CO_2_), complete saturation of gross O_2_ production was observed at around 500 µmol m^-2^ s^-1^ under 2 % CO_2_ (HL-CO_2_). Thus, 500 µmol m^-2^ s^-1^ was selected for HL as a light intensity that saturated photosynthesis under all conditions. Maximum gross O_2_ production was twice as high for HL-CO_2_ as for HL-Amb cells (Supplementary Fig. S1), showing that acclimation to conditions leading to maximum growth rates involved higher photosynthetic electron flow capacity. Under continuous light, gross O_2_ production was halved in ambient CO_2_ compared to 2 % CO_2_ and growth rates were reduced by 15 % under LL and 70 % under HL (Fig. 1, A and B), confirming that cells were CO_2_-restricted even under LL, as previously reported (Zuliani et al., 2024). With 10-min light fluctuations (FL) between HL (FL_HL_) and LL (FL_LL_), cell growth was half under ambient CO_2_ (FL-Amb) compared to 2 % CO_2_ (FL-CO_2_) (Fig. 1A). Aside from the CO_2_ effect, cell density increased faster under FL than under LL, but slower than under HL. Gross O_2_ production during FL_HL_-Amb reached the same rate as HL-Amb, but during FL_HL_-CO_2_ was less than for HL-CO_2_, indicating differential acclimation to HL due to C_i_ availability. Since photosynthetic and light-enhanced dark respiratory (LEDR) rates generally correlated across all growth conditions (Fig. 1, B and C), photosynthetic flux determined mitochondrial flux, at least up to light saturation. However, deviations from this trend occurred in cultures acclimated to conditions approaching and beyond light saturation (i.e. HL and FL_HL_ cells), whereby respiratory flux did not saturate (Supplementary Fig. S1).

**Figure 1.**
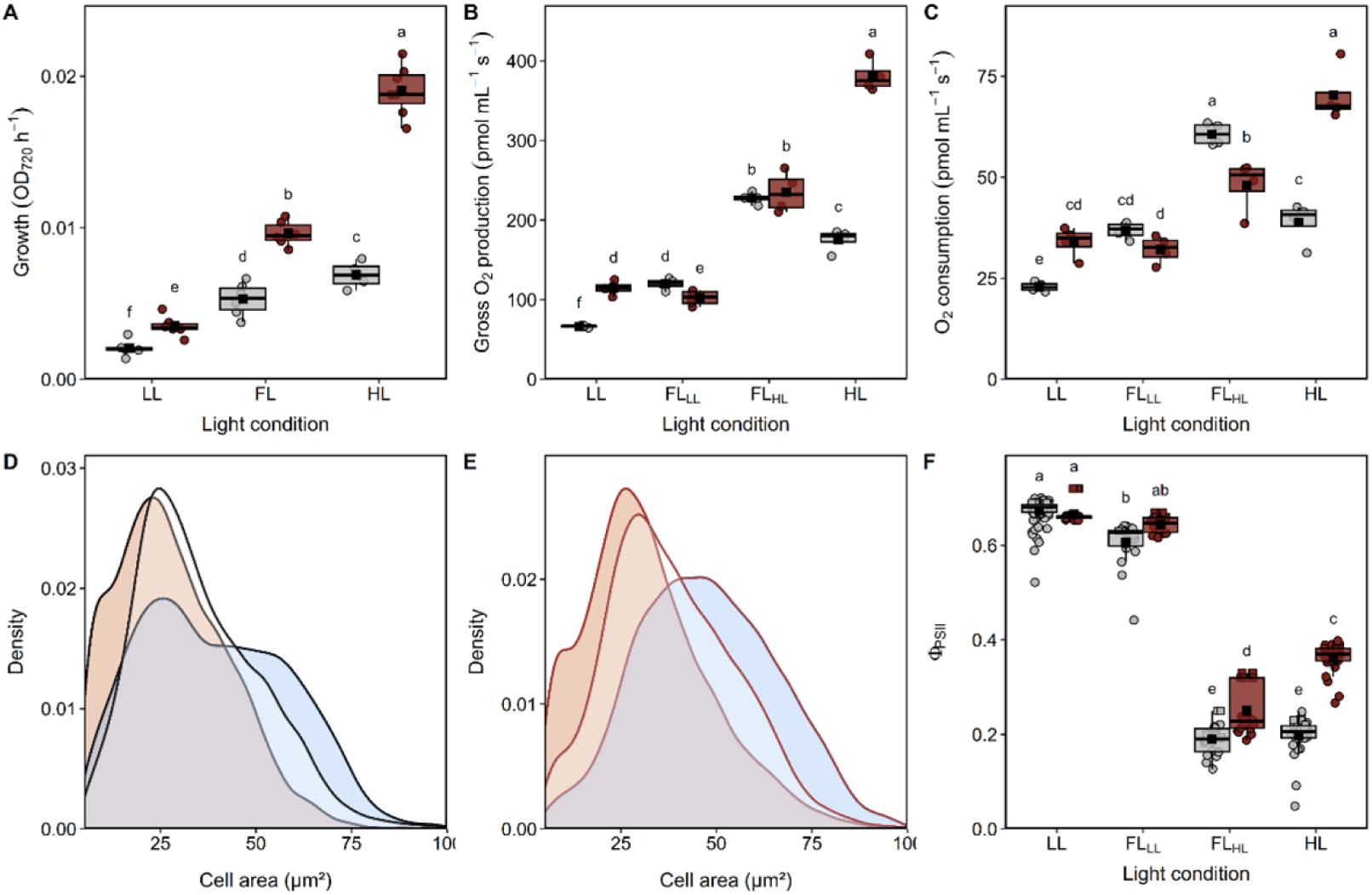
Influence of light regime and CO_2_ concentration on the physiological characteristics of *Chlamydomonas reinhardtii*. Cells were grown in 0.04% (Amb, grey) or 2 % (red) CO_2_, under continuous low light (LL, 50 μmol m^−2^ s^−1^), continuous high light (HL, 500 μmol m^−2^ s^−1^) or fluctuating light (FL), with 10 min duty-cycle switching between LL (FL_LL_) and HL (FL_HL_). (A) Culture growth rates from increases in OD_720_ measured every 5 min during the exponential growth phase from 4-7 independent cultures (*n*= 4-7). (B) Gross oxygen production at steady state in the light under corresponding growth conditions, and (C) oxygen consumption, 30 s after light was turned off, as an indicator of light-enhanced dark respiration (LEDR). Oxygen flux values were corrected based on the OD_720_ (*n*= 4). (D, E) Distribution of cell area of (D) Amb and (E) CO_2_ cells, under LL (blue), FL (white) and HL (red). (F) Relative PSII quantum yields (Φ_PSII_) of cells, in two independent cultivation rounds as measured every 5 min with the internal sensor of the photobioreactor (dots), or externally in an aliquot (squares, *n*= 3). Box plots from panel A-C and F show medians and the 25^th^ and 75^th^ percentiles, dots represent individual data points with dots outside 1.5 × interquartile ranges corresponding to outliers, and black square correspond to the means. Differing letters indicate significant differences (FDR adjusted *p*-value < 0.05).

The median cell area was influenced by growth conditions, shifting to a smaller area with increasing light intensity (Fig. 1, D and E). Additionally, cell area distribution was skewed to larger sizes under LL, and to a lesser extent under FL (Fig. 1, D and E), showing such cells attained a larger area before dividing. Further to indicating a light-dependent speed of the cell cycle, this also showed that using cell number for normalization would not reflect differences in biomass.

Maximum PSII quantum yield (F_v_/F_m_) of HL-CO_2_ and FL-CO_2_ cells was >0.70, and for HL-Amb and FL-Amb cells <0.65 (Supplementary Fig. S2). Hence, under given CO_2_ conditions, F_v_/F_m_ of FL cells was the same as HL cells, which is noteworthy given the distinct growth rates observed under elevated CO₂ conditions. Lower F_v_/F_m_ is a sign that Amb cells had suffered slightly more from PSII photoinhibition. As expected, higher F_v_/F_m_ values were accompanied by higher relative PSII quantum yields (Φ_PSII_) for HL-CO_2_ and FL-CO_2_ compared to their respective Amb cells (Fig. 1F). However, Φ_PSII_ values under FL_HL_-CO_2_ were lower than under HL-CO_2_, indicating a stress on PSII induced by FL. Regarding electron transport rates (ETR), dark-induced reduction kinetics (DIRK) of PSI under HL indicated highest overall electron flux for HL cells and lowest for LL cells (Supplementary Fig. S3), in agreement with gross O_2_ production (Fig. 1B).

Altogether, the physiology of FL-Amb cells resembled HL-Amb cells, whereas FL-CO_2_ cells were more intermediate between LL-CO_2_ and HL-CO_2_ cells. Growth rates and oxygen flux both showed HL-Amb cells were less able to use light energy than HL-CO_2_ cells. Therefore, C_i_ limitation prevented acclimation to HL for achieving maximum growth, rendering Amb-HL and Amb-FL cells during FL_HL_ to HL stress.

### Acclimation to FL involved a partial induction of the photoprotective strategies observed under HL

A total of 1760 proteins were detected using liquid chromatography tandem mass spectrometry (LC-MS/MS) based proteomics in at least one of the six growth conditions, out of which 646 could be quantified in all samples (Supplementary Tables S1 and S2). The light regime exerted the most pronounced influence on the proteome, whereas C_i_ availability showed a more pronounced effect under FL and HL than LL (Supplementary Fig. S4). Availability of C_i_ affected the accumulation of 334 proteins in at least one of the light conditions [false discovery rate (FDR)-adjusted *p*-value < 0.05], out of which 12 consistently showed higher abundance under Amb conditions under all light regimes and another 30 under both FL and HL (*p*-value <0.05, fold change > 1.5; Supplementary Fig. S5, Supplementary Table S3). 9 of those 42 proteins are well-known elements of the CCM whereas two are involved in photorespiration (GLYK1 and GCSP1). Levels of other putative proteins involved in photorespiration, including DLD1, SHMT1 and HPR1, were only higher under FL-Amb compared to FL-CO_2_, indicating an upregulation of this pathway by FL as predicted in plants (Fu and Walker, 2023). Instead, the accumulation of proteins associated to copper and iron deficiency under elevated C_i_ conditions (FOX1, FEA1, FTR1, CHL27A) suggest that the availability of trace elements may become a limiting factor when CO_2_ availability is not restricting growth.

The light conditions caused the differential accumulation of 505 proteins (Supplementary Fig. S6; Supplementary Table S1 and S2). Among these, protein levels of many PSII/PSI subunits, as well as the PSI antenna (LHCA), were relatively less abundant under HL than LL, with typically intermediate levels under FL (Supplementary Fig. S6; Fig. 2A). Regarding the PSII antenna (LHCB), LHCB4 followed the trend for LHCA, while LHCBM3/6 and LHCB5, as well as the NPQ-related LHCSR3, all accumulated most under FL (Fig. 2A). Adjustments in the photosynthetic apparatus, especially with regards to PS and LHC accumulation, have been described as part of the long-term photo-acclimation strategies onset under continuous light (Polukhina et al., 2016). The decrease in photosynthetic complexes per cell can be promoted by the higher cell division frequency under HL (Durnford et al., 2003; Takahashi, 2018). Less light harvesting under HL can enhance light use efficiency with a lower reliance on dissipation mechanisms, while preventing over-reduction of the PETS. In contrast with previous findings (Polukhina et al., 2016), the lesser PS and LHC accumulation was however mostly independent of C_i_ availability, with the notable exception of LHCSR3. Although LHCSR3 expression was only detected in HL-Amb and FL-Amb cells with western blot, LHCSR1 was also detected in HL-CO_2_ and FL-CO_2_ cells (Fig. 2B). *LHCSR* genes are HL-regulated, but *LHCSR3* and *LHCSR1* are under different regulatory pathways (Maruyama et al., 2014). As previously reported (Polukhina et al., 2016; Zuliani et al., 2024), LHCSR3 levels inversely relate with intracellular CO_2_ concentration due to its up-regulation by CIA5 / CCM1, master regulator of the CCM, which does not influence *LHCSR1* expression (Miura et al., 2004; Ruiz-Sola et al., 2023). LHCSR3 typically contributes most to qE-type NPQ (Peers et al., 2009) or protection against PSII and PSI photoinhibition under HL and FL, respectively (Roach, 2020; Roach et al., 2020). Here, chlorophyll fluorescence quenching analyses confirmed the presence of LHCSR3 was necessary for attaining high qE-type NPQ (Fig. 2C). Thus, in agreement with physiological parameters (Fig. 1), HL-Amb and FL_HL_-Amb cells were exposed to light stress and were protecting PS from photoinhibition with NPQ.

**Figure 2.**
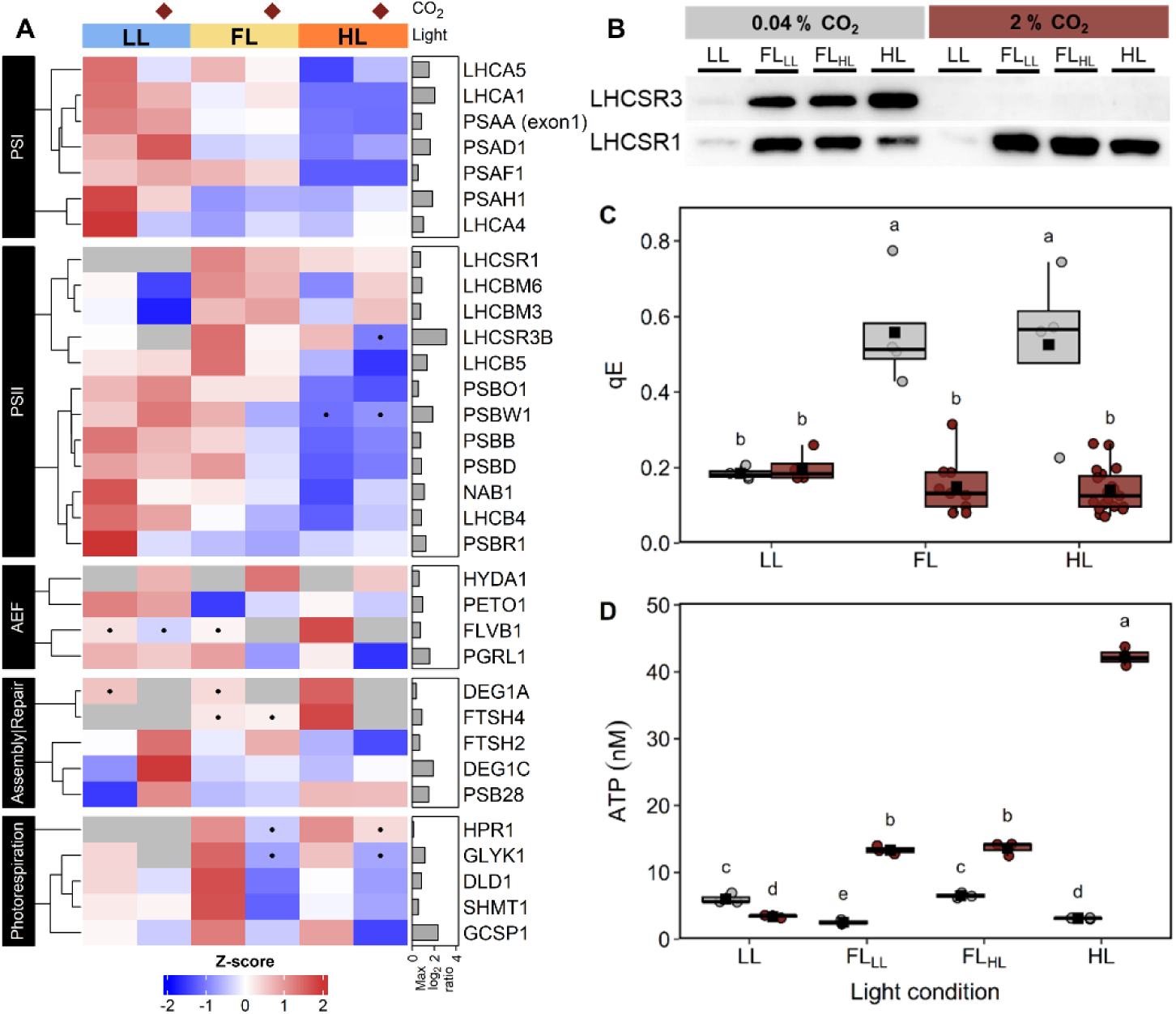
Influence of light regime and CO_2_ concentrations on proteins involved in photosynthesis, high-energy type non-photochemical quenching (qE) and ATP concentration. Cells were grown under 0.04% (grey) or 2 % (red) CO_2_, under continuous low light (LL, 50 μmol m^−2^ s^−1^), continuous high light (HL, 500 μmol m^−2^ s^−1^) or fluctuating light (FL), with 10 min duty-cycle switching between LL (FL_LL_) and HL (FL_HL_). (A) Relative abundances for photosynthetic proteins showing differential accumulation in at least one of the pairwise comparisons (FDR adjusted *p-*value < 0.05) as determined using limma. Z-scores of mean protein relative abundances (*n*= 3) are shown on a color scale from blue (less) to red (more) across growth conditions. Diamonds (top) indicate 2 % CO_2_ cultures and grey bars (right) denote min-max log_2_ fold differences. Protein annotations are provided in Supplementary Table S1. (B) LHCSR1 and LHCSR3 protein abundance, as detected by western blot. (C) High energy state non-photochemical quenching (qE) under HL. (D) ATP concentrations relative to culture volume after OD_720_-based correction. Box plots in C and D show medians and the 25^th^ and 75^th^ percentiles, dots represent individual data points with dots outside 1.5 × interquartile ranges corresponding to outliers, and black squares correspond to the means (*n*= 3). Different letters denote significant differences (FDR adjusted *p*-value > 0.05).

Rates of PSII photoinhibition in the absence of repair under growth light treatments were fastest under HL and only slightly faster under HL-Amb than HL-CO_2_, with no CO_2_ effect detected under LL or FL (Supplementary Fig. S7). Therefore, the lower F_v_/F_m_ (Supplementary Fig. S3) or Φ_PSII_ (Fig. 1F) under HL-Amb and FL_HL_-Amb compared to HL-CO_2_ and FL_HL_-CO_2_, respectively, was presumably due to differences in repair rates. DEG and FTSH proteases, potentially related to PSII repair (Malnoe et al., 2014; Theis et al., 2019), showed differential accumulation under the different conditions. DEG1A and FTSH4 mostly accumulated in Amb cells, with highest levels under HL indicating an association with stress, whereas DEG1C and FTSH2 were more abundant in LL-CO_2_ cells. The reason for this is unclear, but it does show that C_i_ availability influences the regulation of proteases. Nonetheless, the repair of PSII has a major energetic cost for cells, consuming vast amounts of ATP (Raven, 2011). Hence, C_i_ availability presumably enhanced photosynthetic efficiency under FL and HL by promoting PSII repair, rather than preventing photodamage, which points to a higher ATP availability.

### The cellular energy balance under FL is CO2 dependent

Growth under 2 % CO_2_ led to substantially higher ATP concentrations under FL and HL than under ambient CO_2_ (Fig 2D). Instead, HL-Amb cells accumulated the least ATP, which may reflect both the high demand for ATP in repair processes, and the lower photosynthetic efficiency resulting from photoinhibition and enhanced energy dissipation through NPQ. The highest ATP concentration measured in HL-CO_2_ cells was also reflected in their growth rates. Indeed, as cell division comes with a high energetic cost, growth rates are highly dependent upon the cellular energetic balance, and cell division may represent an effective sink for excess reductants (Davis et al., 2013). The difference in ATP concentration between cells grown under LL with or without CO_2_ supplementation may reflect differences in light acclimation strategies, leading to a higher investment into growth under LL-CO_2_. Cells grown under FL showed intermediate ATP concentrations as compared to the corresponding continuous light conditions. The ATP concentration in cells grown under FL-Amb was substantially higher under FL_HL_ than under FL_LL_, indicating a light phase dependent regulation of the cellular energetic balance. In contrast, cells grown under FL-CO_2_ had at least twice more ATP, whereby ATP concentration was comparable at the end of both light phases, suggesting that compensatory mechanisms were induced to maintain the cellular energy balance. Such mechanisms possibly relied on transient metabolite pools which accumulation was C dependent.

Cells use alternative electron flows (AEF) to protect against photoinhibition and to balance NADPH:ATP ratios for metabolism. Linear electron flow (LEF) produces insufficient ATP relative to NADPH for photosynthetic carbon fixation. Hence, the provision of additional ATP depends on a number of AEF, including CEF around PSI, involving a complex containing anaerobiosis Response 1 (ANR1), Proton Gradient Regulation 5 Like 1 (PGRL1) and the transmembrane thylakoid phosphoprotein (PETO) (Takahashi et al., 2016). Whereas ANR1 was only marginally higher in FL-Amb than LL-Amb (Supplementary Tables S1 & S2), both PGRL1 and PETO1 were most abundant in cells acclimated to LL (Fig. 2A), which agreed with the highest PSI:PSII ETR ratio for LL-acclimated cells (Supplementary Fig. S8). Regarding PCEF, net oxidation from ca. 200 ms during the saturating pulse is putatively FLV activity, due to its absence in *FLV* knockout mutants (Pfleger et al., 2024). Here, P700 maintained a more oxidized state from ca. 200 ms in Amb compared to CO_2_ acclimated cells (Supplementary Fig. S9), agreeing with the higher FLVB1 levels in the proteomic data (Fig. 2A). Moreover, under FL-CO_2_, P700 was hardly oxidized during measurements, while under FL-Amb P700 was less oxidized than under HL-Amb, indicating that FLV activity was lower under FL despite that FL conditions can be intolerable for *flvb1* with a high light phase of 1 min (Chaux et al., 2017). A higher PCEF under HL may help prevent an over-acidification of the lumen due to CEF (Kramer et al., 1999). Overall, CEF was likely most important for ATP production under LL, while PCEF contributed least under FL, although enhanced Mehler reaction due to low FLV activity cannot be excluded (Pfleger et al., 2024).

### The activation of CCM is light inducible and results in enhanced energy trade-offs between cellular organelles

The carbon concentrating mechanism (CCM) is induced by low CO_2_ levels (Mackinder et al., 2017). In our study, growth rates revealed C_i_ limitation in Amb cells under all light treatments, which was associated with the increased abundance of important CCM components (Fig. 3), including transporters (the bicarbonate channels LCI1 and NAR1B/LCIA, the plasma membrane H^+^ ATPase PMA3), carbonic anhydrases (the periplasmic CAH 1 and 2, thylakoid lumen-localized CAH3, mitochondrial CAH4 and three ß-like carbonic anhydrase proteins LCIB, LCIC and LCID) and pyrenoidal linker proteins (RBMP2, EPYC1). Acclimation to limiting C_i_ is accompanied by changes in the subcellular organisation in *C. reinhardtii*, including the formation of the pyrenoid and the allocation of starch to the establishment of a starch sheath surrounding it (Kuchitsu et al., 1988; Izumo et al., 2011). The starch sheath is considered to restrict CO_2_ diffusion out of the pyrenoid, hence reducing the energetic cost of operating the CCM, and may also contribute in preventing RuBP oxygenation, by segregating RubisCO away from sites of O_2_ evolution (Toyokawa et al., 2020; He et al., 2023). Indeed, LL-Amb and FL-Amb cells contained higher starch concentrations than LL-CO_2_ or FL-CO_2_, reflecting the formation of pyrenoidal starch (Fig. 4). GBSS1A (STA2), a granule bound starch synthase presumably contributing to the formation of the starch sheath, was found up-accumulated in LL-A and FL-A cells. *GBSS1A* expression is positively regulated by the CCM regulator CIA5 (He et al., 2023). Unlike the strong influence of CO_2_ on the CCM and starch contents, we found no effects of growth conditions on any proteins directly assigned to RubisCO activity (e.g. RBCL, RAF1, CPN20/23; Supplementary table S1). Cytoskeletal regulation of mitochondrial movements under limiting C_i_ conditions lead to repositioning of the mitochondria from a central position within the cup of the chloroplast to the periphery of the cell. (Geraghty and Spalding, 1996; Findinier et al., 2024). The mitochondrial CAH4 and the putative HCO_3_ ^-^ transporter CCP1, which expression were shown to be induced under very low CO_2_ conditions (Findinier et al., 2024)(< 0.02% CO_2_, Findinier et al. 2024), were both strongly accumulated under FL-Amb and HL-Amb. Those proteins were proposed as part of a mechanism enabling the recapture of CO_2_, especially that generated by respiratory processes or leaking out from the chloroplast, to increase the cell’s affinity for C_i_ (Rai et al., 2021). Their higher accumulation under conditions promoting CO_2_ consumption, together with that of other CCM related proteins (LCI1, LCI9, LCID, LCI19), could therefore be a response to the increasing C_i_ demand under higher light intensities and / or to increased reliance on respiration, as previously suggested (Mettler et al., 2014).

**Figure 3.**
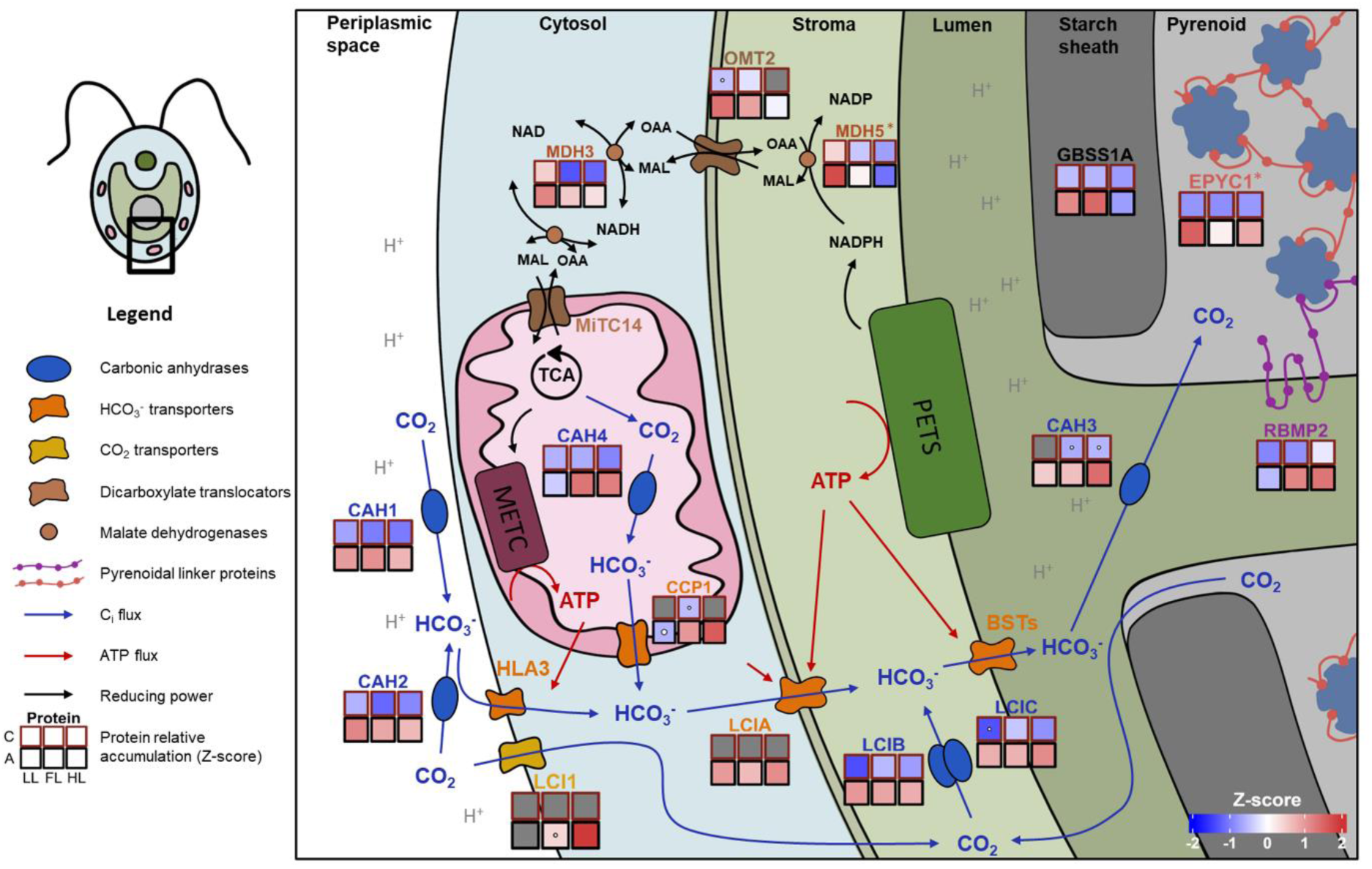
Interaction between the carbon concentrating mechanism (CCM) and the malate valve in response to different light and CO_2_ conditions. Cells were grown in 0.04% (A) or 2 % (C) CO_2_, under continuous low light (LL, 50 μmol m^−2^ s^−1^), continuous high light (HL, 500 μmol m^−2^ s^−1^) or fluctuating light (FL), with 10 min duty-cycle switching between LL and HL. CCM components includes HCO_3_^-^ transporters [orange, HLA3, LCIA, bestrophins (BSTs), CCP1], CO_2_ transporter (yellow, LCI1), carbonic anhydrases (blue, CAH1, CAH2, CAH3, CAH4) and ß-like carbonic anhydrases (blue, LCIB, LCIC), which in conjunction convert CO_2_ to HCO_3_^-^ and then transport it towards the pyrenoid for carbon fixation. The translocation of protons (H^+^) leads to the acidification of the lumen and periplasm, which increases the activity of the CAHs. The malate shuttle components (brown) include the carboxylate transporter OMT2 and malate dehydrogenases (MDH3/5), which can shuttle reducing equivalents across cellular compartments. GBSS1A functions in starch synthesis in the sheath. The pyrenoidal linker proteins [EPYC1 (red) and RBMP2 (purple)] are linked to Rubisco and are involved in the pyrenoid assembly under low CO_2_ conditions. Z-scores of mean protein relative abundances (*n*= 3) are shown in each square on a color scale from blue (less) to red (more) (grey = not detected; white dots = not detected in one or two of the replicates) across growth conditions (key left). Only proteins differentially accumulated in at least one of the pairwise comparisons are shown (FDR adjusted *p*-value< 0.05; except for protein marked with an asterisks for which only the non-adjusted *p*-value was below 0.05). The figure was inspired by Mackinder et al, 2017, Burlacot & Peltier, 2023 and Burlacot et al., 2022. Abbreviations: MAL, malate; METC, mitochondrial electron transfer chain; OAA, oxaloacetate; PETS, photosynthetic electron transfer chain. Refer to Supplementary Table S1 and S2 for information about proteins.

**Figure 4.**
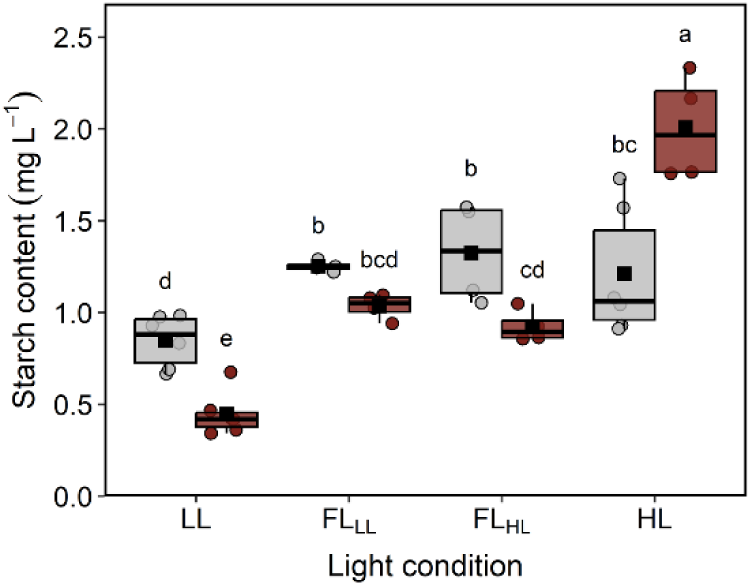
Influence of light regime and CO_2_ concentrations on the starch content. Cells were grown under 0.04% (grey) or 2 % (red) CO_2_, under continuous low light (LL, 50 μmol m^−2^ s^−1^), continuous high light (HL, 500 μmol m^−2^ s^−1^) or fluctuating light (FL), with 10 min duty-cycle switching between LL (FL_LL_) and HL (FL_HL_). Amounts are relative to total culture volume after OD_720_-based correction. Box plots from show medians and the 25^th^ and 75^th^ percentiles, dots represent individual data points with dots outside 1.5 × interquartile ranges corresponding to outliers, and black squares correspond to the means (*n*= 4-6). Different letters denote significant differences (FDR adjusted *p*-value > 0.05).

The proximity of the relocated mitochondria to CCM related carbon transporters in the plasma membrane highlight the involvement of the mitochondria in CCM, for example by powering the carbon transport into the cells through the malate shuttle (Burlacot and Peltier, 2023). This is consistent with the latest studies, which indicate that malate shuttle components are responsive to CO_2_ levels and possibly involved in the algal CCM (Dao et al., 2022). Under conditions of limited C_i_ concentration when CCM in algae is active, additional ATP is needed for CO_2_ or bicarbonate transport (1/2 additional ATP molecules/fixed CO_2_) and for pyrenoidal starch sheath formation (1 ATP/Glucose) (Johnson and Alric, 2013). Hence, the energetic cost of the CCM may explain the lower ATP content of cells grown under FL-Amb as compared to FL-CO_2_ conditions. We observed an enhanced response of the malate shuttle components under limiting C_i_ conditions, including the chloroplastic malate dehydrogenase 5 (MDH5), the cytosolic malate dehydrogenase 3 (MDH3) and the putative 2-oxoglutarate/malate antiporter (OMT2), which lends support to the hypothesis that malate shuttles are involved in the energy supply to the CCM. However, the specific MDHs involved in exchanging reducing equivalents between mitochondria and chloroplasts remain unknown (Burlacot et al., 2019). In relation to the various light treatments, an increase in the malate shuttle components (OMT2, MDH5) was observed under FL-Amb and LL conditions. This suggests an enhanced shuttling of reductants from the chloroplasts to the mitochondria, in which the malate shuttle may be important for the aforementioned energy balancing of ATP/NADPH when light is sub-saturating, as described in *Arabidopsis* (Walker et al., 2014; Yokochi et al., 2021). With increasing light, other alternative mechanisms (e.g. FLV) may modulate the ATP/NADPH supply.

### Growth under FL involved coordinated inter-compartmental metabolic processes tuning protein synthesis to the energy balance

Overall, the primary metabolite profiles of cells grown under FL showed substantial variations depending on the light phases, whereby the accumulation of many metabolites at the end of the LL and HL phases mirrored that of cells continuously grown under the corresponding light intensities and CO_2_ levels (Fig. 5, Supplementary Fig. S10, Supplementary Tables S4 and S5). C_i_ availability had an influence on accumulation of primary metabolites, especially under HL and resulted in higher accumulation of most detected metabolites. Under both Amb and CO_2_, an accumulation of Calvin Benson Bassham (CBB) cycle intermediates, namely sedoheptulose-7-phosphate, ribose-5-phosphate, ribulose-5-phosphate and fructose-6-phosphate, was observed during the HL phase of the FL regime followed by a marked decrease within 30 s upon transition to LL (Fig. 5). Hence, fluxes through the CBB cycle rapidly responded to light fluctuations, regardless of C_i_ availability. Differences in CBB intermediate levels did not directly correlate with changes in the proteins involved (Supplementary table S1). Instead, the accumulation of many CBB enzymes to similar or higher concentrations than their substrates may underlie the capacity to sustain higher fluxes under FL_HL_ (Mettler et al., 2014). Under elevated CO_2_, intermediates downstream of carbon fixation shared with the glycolysis / gluconeogenesis pathways, namely glyceraldehyde-3-phosphate (G3P), dihydoxyacetone-phosphate (DHAP), 3-phosphoglycerate and fructose-1,6-biphosphate, and TCA cycle intermediates (succinate, fumarate, malate, citrate) showed a similar pattern but with a slower decline during the LL phase. These increased levels of intermediates reflected higher metabolic fluxes through those pathways during the HL phase, when C assimilation and energy were not limiting. Instead, under Amb conditions, levels of most glycolysis and TCA cycle intermediates remained comparatively lower regardless of the light condition, consistent with a substantial influence of C availability on the fluxes through the primary metabolic pathways.

**Figure 5.**
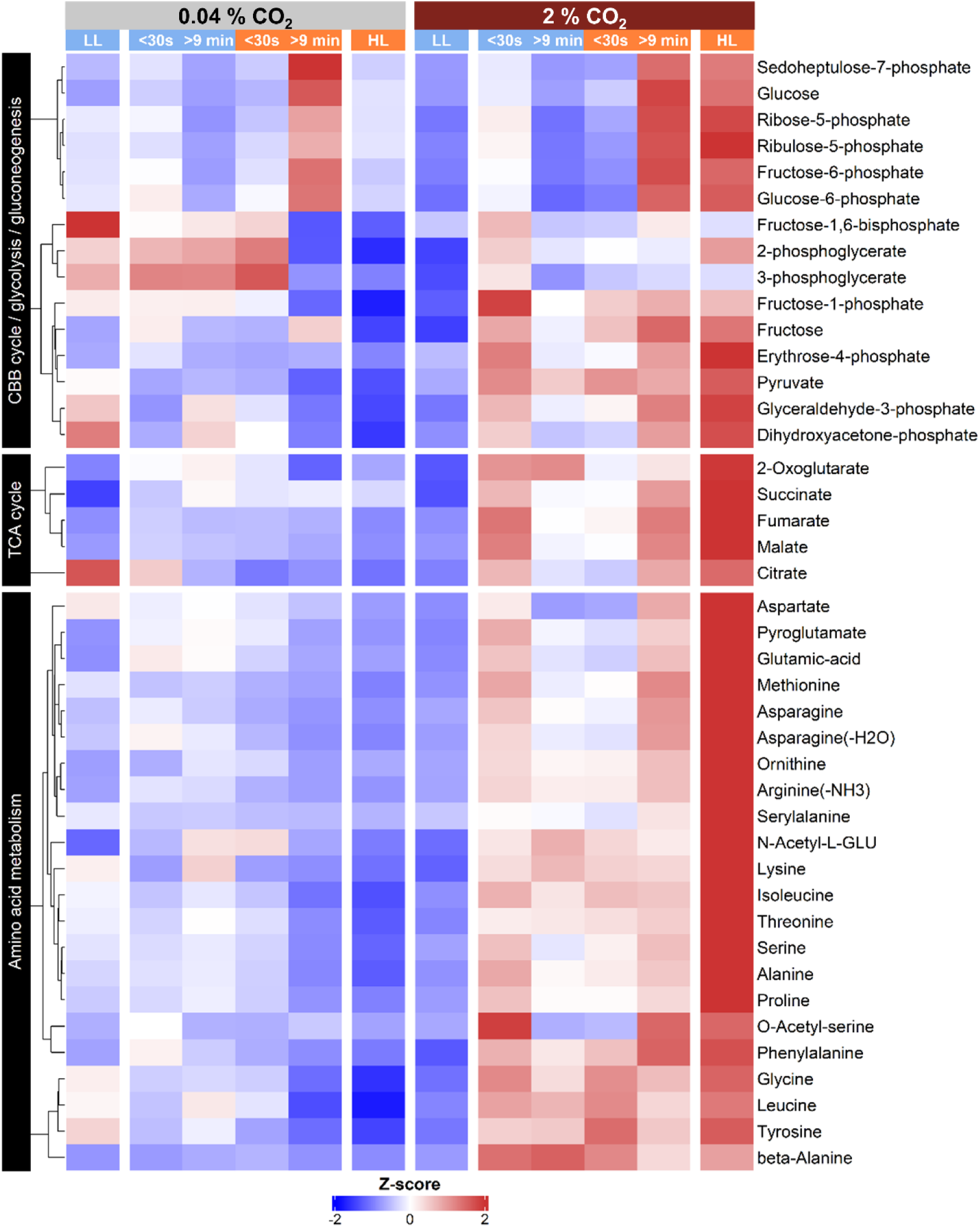
Dynamic response of core metabolic pathways to light fluctuation as influenced by C_i_ availability. Cells were grown under 0.04% (grey) or 2 % (red) CO_2_, under continuous low light (LL, 50 μmol m^−2^ s^−1^), continuous high light (HL, 500 μmol m^−2^ s^−1^) or fluctuating light, with 10 min duty-cycle switching between LL and HL. Metabolites are grouped into the Calvin Benson Bassham (CBB) cycle and glycolysis / gluconeogenesis pathway, the tricarboxylic acid (TCA) cycle and amino acid metabolism. FL cells were harvested at the beginning (< 30 s) and end (>9 min) of each LL and HL phase. Z-scores of mean metabolite relative abundances are shown on a color scale from blue (less) to red (more) across growth conditions (*n*= 5). Only metabolites that showed differential accumulation in at least one pairwise comparison (FDR adjusted *p*-value < 0.05) are shown. Within each group, metabolites were hierarchically clustered using the Euclidean distance and Ward’s clustering method.

Triose-phosphates (G3P & DHAP) produced from the CBB cycle or glycolysis in the chloroplast, are the main photo-assimilates exported to the cytosol by triose phosphate/inorganic phosphate translocators (TPT) for further conversion to pyruvate, which can feed into the TCA cycle to sustain respiration and mitochondrial ATP production (Johnson and Alric, 2013; Moulin, 2023). TPT3, which was detected under all culture conditions, plays a pivotal role in the export of photo-assimilates from the chloroplast and can act as a safety valve preventing excess reducing power from the chloroplast (Huang et al., 2023). This inter-compartmental metabolite shuttling may substantially account for the LEDR, which increased with increasing light intensities (Fig. 1C). In conditions under which the availability of C skeletons issued from the glycolysis or malate shuttle becomes limiting, amino acids may also serve as substrates to sustain ATP production. Taken together with a reduced C assimilation, the higher ATP demand for the CCM under Amb conditions may lead to a faster consumption of available assimilates, especially under LL conditions. Instead, under lower ATP demand (e.g. HL-CO_2_ / FL_HL_-CO_2_), reactions branching from the TCA cycle can divert C skeletons toward multiple anabolic processes, including nitrogen assimilation and amino acid biosynthesis (Sweetlove et al., 2010). Hence, an increased flow through the pathways of energy metabolism, especially the TCA cycle, under elevated CO_2_ may not only contribute to ATP production, but also to amino acid (AA) synthesis, as supported by the clear accumulation observed for most AAs under both HL-CO_2_ and FL-CO_2_ (Fig. 5). Under FL, the balance between cyclic and non-cyclic fluxes through the TCA cycle may be key in tuning ATP production and biomass accumulation depending on light availability. Indeed, ATP concentrations did not change between FL_HL_ and FL_LL_ neither under CO_2_ nor Amb (Fig. 2D). The predominance of non-cyclic flows of TCA cycle intermediates could account for the generally lower LEDR under FL-CO_2_ as compared to FL-Amb cells (Fig. 1C). Increased AA pools can in turn promote protein turnover and *de novo* synthesis. Accordingly, the total protein content substantially increased with increasing light intensities and C_i_ availability (Fig. 6), closely reflecting the increase in growth rates (Fig. 1A). Hence, increased energy and AA availability were consistent with higher protein contents, supporting faster growth, in agreement with previous findings under continuous HL (McKim and Durnford, 2006; Davis et al., 2013).

**Figure 6.**
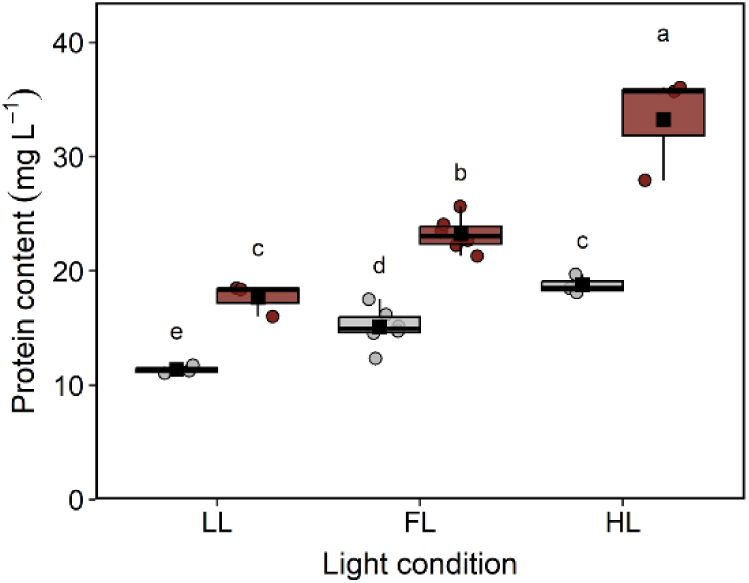
Influence of light regime and CO_2_ concentrations on the protein content. Cells were grown under 0.04% (grey) or 2 % (red) CO_2_, under continuous low light (LL, 50 μmol m^−2^ s^−1^), continuous high light (HL, 500 μmol m^−2^ s^−1^) or fluctuating light (FL), with 10 min duty-cycle switching between LL and HL. Amounts are relative to total culture volume after OD_720_-based correction. Box plots from show medians and the 25^th^ and 75^th^ percentiles, dots represent individual data points with dots outside 1.5 × interquartile ranges corresponding to outliers, and black squares correspond to the means (*n*= 3-6). Different letters denote significant differences (FDR adjusted *p*-value > 0.05).

### Transient photo-assimilate pools serve as energy buffers under FL while excess of Ci under HL results in accumulation of starch reserves

Starch, amongst other metabolites, can provide a transient buffer for C within the first 10-15 minutes of increased light exposure in *C. reinhardtii*. This buffering mechanism may be maintaining the C balance and metabolic stability during light transitions, when light switches from HL to LL (Mettler et al., 2014). Accordingly, the increased C_i_ assimilation under the HL phase of FL led to the transient accumulation of glucose and hexose-phosphates, which appeared to be remobilized to sustain the metabolic activity upon transitioning to LL under both ambient and elevated CO_2_ conditions. A rapid decline in chloroplastic ATP production consecutive to the transition from HL to LL may relieve the allosteric inhibition of phosphofructokinases (PFK) (Johnson and Alric, 2013), especially of PFK2 which accumulated under HL and FL as compared to LL, allowing for a rapid response to changes in cellular energy status. This would in turn enable the remobilization of hexoses and hexose-phosphates through glycolysis, allowing for a sustained overall ATP production through mitochondrial respiration. Such a mechanism could explain the delayed decrease observed for glycolysis and TCA cycle intermediates upon transition from HL to LL. During the HL phase, PFK inhibition due to increased chloroplastic ATP production may instead promote starch synthesis, in line with the higher accumulation observed under FL for several enzymes involved in starch synthesis, including a phosphoglucomutase (PGM1) and PHOB1/STA4. PGM1 catalyzes the first step in directing the end products of photosynthesis towards starch storage and is allosterically up-regulated by light and products of photosynthetic activity, e.g. glucose-1,6-bisphosphate (Sicher and Kremer, 1990). The α-amylases 2 and 3 (AMA2, AMA3) also showed higher accumulation under FL. Hence, the up-accumulation of enzymes involved in starch synthesis and breakdown under FL, together with the transient accumulation of hexoses and hexose phosphates, could indicate that *C. reinhardtii* uses starch as a temporary C storage to meet changing metabolic demands under FL conditions.

When grown under LL or FL, cells accumulated more starch under Amb rather than CO_2_ conditions, whereas the reverse was observed for cells grown under HL (Fig. 4). Under 2 % CO_2_, starch content increased with increasing light intensity, but no differences were found between the LL and HL phases of FL, which suggests that hexoses and hexose-phosphates may be more important in buffering the energetic pathways under the FL conditions applied. When growth under HL is not restricted by C limitation, an overflow of C skeletons produced through photosynthesis results in storage starch accumulation in the stroma, as observed previously (Polukhina et al., 2016). The starch synthases, SSS3A (STA3), showed its highest accumulation under HL-CO_2_ and FL-CO_2_, conditions under which it likely promoted stromal starch accumulation, which can then function as a sink and source of sugar for supporting metabolism as required.

Lipids can represent an alternative form of storage sink for reduced C. However, their accumulation is only favoured under specific conditions, including nitrogen deprivation or mixotrophic conditions (Fan et al., 2012). Accordingly, total fatty acids (FAs) only showed limited variations in response to changes in light intensity and C_i_ availability (Supplementary Fig. S11). Under Amb, content of the monounsaturated FA oleic (C18:1Δ^9^) and palmitoleic acid (C16:1Δ^7^) substantially decreased with increasing light intensity, whereas they slightly accumulated under HL-CO_2_.In contrast, a slight increase was observed for most of the other FAs with increasing light intensity under CO_2_, overall indicating that some of the excess C skeletons were incorporated into lipids.

### Conclusion - Growth limitation due to energy trade offs

The PETS immediately responded to changes in light intensity, leading to rapid metabolic adjustments under FL, which were influenced by CO_2_ availability and linked to growth performance. Growth limitation by C_i_ restriction increased under HL and reflected the increased total C demand. The growth performance under FL was hampered by light limitation during FL_LL_ phases and by excess energy absorption under FL_HL_ phases, which either contributed to enhanced C_i_ fixation or led to enhanced energy dissipation depending on C_i_ availability.

Light and CO_2_ availability determine ATP sinks. Under C_i_ restriction, induction of the CCM increased with light intensity and comes with an energetic cost for its establishment (CCM gene expression, pyrenoid starch sheath) and operation (transporters, proton pumps). Under HL and FL_HL_, ATP for the PSII repair is needed, due to higher photoinihibition, and the lower Φ_PSII_ and higher NPQ under FL-Amb and HL-Amb relative to 2 % CO_2_ condition is likely reflecting the lower ATP concentration in the cells. Overall, the costs imposed by the CCM, photoprotection and PSII repair, represent energy trade-offs required for cell survival at the cost of growth under suboptimal conditions.

The tight regulation of metabolism, especially through energetic exchanges between cellular compartments, facilitated by malate shuttles and export of photo-assimilates from the chloroplast, allows for a high metabolic flexibility. Whereas rapid changes were observed for primary metabolites directly connected to photosynthesis in response to FL, the delayed variations observed for downstream intermediates exchanged between compartments likely reflected the importance of transient reserve accumulation acting as energy and C buffers. The resulting cellular energy balance drives C allocation and is crucial for growth regulation especially depending on the environmental conditions. Overall, elevated CO_2_ favors the energy balance under HL and FL_HL_ which promotes investment of C skeletons into biomass production, such as storage starch and protein synthesis, and enables rapid cell divisions. As a result, C_i_ availability is a key determinant of growth performance under FL.

## Methods

### Cultivation

The *Chlamydomonas reinhardtii* wild-type (WT) strain CC-4533 was kept on 1.5 % agar supplemented with tris-acetate-phosphate (TAP) medium in a growth chamber (at 25 °C, 25 µmol m^-2^ s^-1^). For experiments, strains were pre-cultured in 1 L Schott bottles containing photoautotrophic Kropat’s medium (Kropat et al., 2011) and HEPES buffer, which was adjusted to a pH of 7.5 with potassium hydroxide. Cultures were shaken in a growth chamber at 25 °C and 100 rpm under continuous white light (100 µmol m^-2^ s^-1^) and bubbled with sterile pre-humidified air prior to starting treatments. Experimental cultures were then transferred in the early exponential phase to 1 L flat-panel photobioreactors FMT150 (PSI, Drásov, Czech Republic) at 25 °C and aerated with constant sterile and humidified air bubbling at an airflow of 0.5 L/min (CO_2_ = 0.04 %, O_2_ = 21 %) or elevated CO_2_ (CO_2_ = 2%, O_2_ = 21 %) and maintained under continuous HL (500 µmol m^-2^ s^-1^), continuous LL (50 µmol m^-2^ s^-1^) and FL (10 min duty cycle between HL and LL) for at least 48 h. Several cultivation rounds were conducted. Growth was monitored using optical density at 720 nm (OD_720_), as measured every 5 min by the FMT150 photobioreactor. To minimize self-shading, all experiments were performed with cultures early in exponential growth at an OD_720_ of up to 0.3 that corresponded to a chlorophyll concentration of ca. 2.5 µg mL^-1^.

### Cell size evaluation

Cell counts were determined microscopically in a Fuchs-Rosenthal counting chamber (cell depth: 0.2 mm, total cell area: 16mm^2^) using a fully motorized Olympus (BX53) microscope (Evident, Waltham, MA, USA) at 2000-fold magnification. Deep-learning technology applied through the Olympus cellSens imaging software (v3.2, Evident) was used for automated identification and measurement of size and area of cells grown under each growth condition.

### High-resolution O2 flux measurement

Net O_2_ production and consumption were measured using a NextGen-O2k (Oroboros Instruments, Innsbruck, Austria) connected to a PB-Module, consisting of two 2-mL chambers each connected to a Clark-type polarographic O_2_ sensor (POS). All experiments were conducted under constant stirring (750 rpm) at 25 °C. Data were collected every 2 s. As the experimental chamber was a closed system during measurements, measurements were made in 5 mM NaHCO_3_ to ensure that photosynthesis was saturated with CO_2_. Instrument calibration was performed with cell-free medium according to (Gnaiger, 2001, 2008). The volume-specific O_2_ flux (O_2_ flux per volume) was calculated as the time derivative of the O_2_ concentration corrected for instrumental background using the DatLab software (v8.0.3, Oroboros Instruments). The reported O_2_ fluxes correspond to the median of 5-30 data points after signal stabilization, which were corrected based on the OD_720_. To investigate the light response, cells were exposed to 50, 250, 500 and 750 µmol m^-2^ s^-1^. The protocol was automated with cycles for each light intensity starting with 5 min in the dark, followed by 5 min in the light. The peak of O_2_ consumption that occurred 30 s after switching off the light was used as a measure of LEDR. The subtraction of LEDR from net O_2_ production allowed to calculate the gross O_2_ production.

### Metabolite profiling

For metabolite profiling, cultures were grown to an OD_720_ of 0.3 ± 0.02 before harvesting. For FL, cells were harvested < 0.5 min and > 9 min after a change in light intensity, whereas cells grown under continuous HL or LL were harvested once. In total, five samples were taken from each growth condition (*n*=5). Metabolite profiling was conducted according to (Lee and Fiehn, 2008; Fiehn, 2016), with adaptations of the protocol. First, 25 mL of culture was harvested with a 50 mL syringe and immediately quenched in 25 mL 60 % methanol pre-cooled to −50 °C. After centrifugation (2000 *g*, 3 min, −20 °C) the supernatant was discarded and cells were flash-frozen in liquid N_2_ and kept at −80 °C until extraction. Metabolites were extracted in methanol:chloroform:water (10:3:1, v:v:v), supplemented with 21.26 µM ^13^C_6_-sorbitol (Campro Scientific GmbH, Berlin, Germany) and 25 µM ^13^C_6_, ^15^N-valine as internal standards and degassed with nitrogen for 5 min. After adding 500 µL of extraction solvent pre-cooled to −20 °C, samples were stirred with two 3 mm glass beads at 4°C and 800 rpm for 10 min, on a Compact Digital Micro plate shaker (Thermo Fisher Scientific, Waltham, MA, USA) before transfer to 2 mL reaction tubes. Samples were kept in racks precooled to −20 °C during the entire extraction. Insoluble material was pelleted by centrifugation (20 000 *g*, 5 min, 4 °C) and kept for starch and protein content assessments. The supernatant was combined with 200 µL of water, vortexed for 15 s and centrifuged (20 000 *g*, 2 min, 4 °C) to induce phase separation of polar (methanol/water) and lipophilic phases (chloroform). 100 µL aliquots of the polar phase was collected for metabolite profiling and dried for 3 h in a vacuum centrifuge (SpeedVac SPD111V P2, Thermo Fischer Scientific). Metabolites were chemically derivatized in two steps, first by incubation with 10 µL of methoxyamine hydrochloride at 20 mg mL^-1^ in pyridine at 28 °C and 600 rpm for 90 min, followed by an incubation with 90 µL of N-methyl-N-trimethylsilyl-trifluoroacetamide (MSTFA) at 37 °C and 600 rpm for 30 min, in both cases using a thermomixer (Eppendorf Austria GmbH, Vienna, Austria).

A 1 µL aliquot of the resulting sample was injected in the split-splitless inlet of a 7890B gas chromatograph (Agilent, Santa-Clara, CA, USA), operated at 250 °C in splitless mode with a carrier flow of helium at 1 mL min^−1^. Analytes were separated on a 30 m Rxi-5Sil MS column with a 10 m Integra-Guard pre-column (Restek, Bellefonte, PA, USA) using an oven temperature ramp starting at 70 °C for 7 min, followed by an increase in 10 °C min^−1^ up to 325 °C, which was held for 7 min. Mass spectra were acquired after 9 min solvent delay, using a Pegasus BT time of flight mass spectrometer (LECO Corporation, St. Joseph, MI, USA), scanning from 50 to 550 m/z at a frequency of 15 spectra s^−1^, with the transfer line and ion source temperatures, respectively, set to 290 and 250 °C. Between consecutive injections, the 10-μL syringe was washed four times each with hexane and with ethyl acetate. A mix of alkanes dissolved at 2 mg L-1 in hexane was injected in the middle of the queue to allow for the conversion of retention times into Kováts’ alkane-based retention indices (Kovats, 1958). Data acquisition and review was conducted using the ChromaTof software (version 5.56.57, LECO) in combination with the National Institute of Standards and Technology (NIST, 2020 release), Golm and Fiehn mass spectral libraries for compound identification (Kopka et al., 2005; Kind et al., 2009), based on matches of both spectral data and retention indices. Relative metabolites abundances were calculated by normalizing peak areas for compound-specific ions to those of the internal standards and to the OD_720_ measured upon sampling.

### Fatty acid profiling

Total fatty acids analysis was conducted through derivatization to fatty acid methyl esters (FAMEs) as previously described (Pichrtová et al., 2016). Briefly, 50 µL of the chloroform phase was fully evaporated under nitrogen using a Reacti-Vap evaporator (Thermo Fisher Scientific) and resuspended in 500 µL of methanol:toluene:sulphuric acid 40:12:1 (v:v:v) containing 0.01 % (w:v) butylated hydroxytoluene. After 90 min at 80 °C with constant agitation, 160 µL of hexane and 600 µL of ultrapure water containing 0.9 % NaCl (w:v) were added to each sample, followed by vigorous mixing and centrifugation (6000 *g,* 6 min). One µL of the resulting supernatant was then injected in the split-splitless inlet of a Trace 1300 gas chromatograph (Thermo Fisher Scientific), operated at 225 °C in split mode with a carrier flow of helium of 1.5 mL min^−1^ and a split flow set to 15 mL min^−1^. FAMEs were separated on a 30 m BPX70 column (Trajan Scientific, Ringwood, Australia) and detected using a TSQ 8000 triple quadrupole mass spectrometer (Thermo Fischer Scientific) operated in full scan mode (50– 450 m/z) at 5 spectra s^-1^ after 3.5 min of delay. The GC oven temperature was first set to 70 °C for 12 s, then increased by 10 °C min^−1^ up to 180 °C, and finally by 15 °C min^−1^ up to 230 °C, which was held for 5 min. The ion source and transfer line temperatures were set to 250 °C and 235 °C, respectively. Data acquisition and review was conducted using the Xcalibur software (v4.5, Thermo Fischer Scientific), following the above-described approach to assess fatty acid relative abundances.

### Proteomics using LC-MS/MS

Cells were harvested at an OD_720_ of 0.23 ± 0.07 by centrifugation (2000 *g*, 5 min) at 4 °C. After discarding the supernatant, cells were flash-frozen in liquid N_2_, freeze dried and stored at −80 °C. Cells were then ground for 2 min in a ball mill using 2 mm steel balls and proteins were extracted, pre-fractionated by 1D SDS-PAGE (using 40 μg of total protein), trypsin digested and desalted using a C18 spec plate as previously described (Chaturvedi et al., 2013; Ghatak et al., 2021). Prior to mass spectrometric measurement, the tryptic peptide pellets were dissolved in 4% (v/v) acetonitrile with 0.1% (v/v) formic acid. For each sample, one µg of purified peptides was loaded onto a C18 reverse-phase analytical column (EASY-Spray 50 cm, 2 µm particle size, Thermo Fisher Scientific) using a Dionex Ultimate 3000 RSLCnano liquid chromatography system (Thermo Fisher Scientific). Separation was achieved using a 140 min gradient method, first ramping from 5% to 50% solvent B (v/v) [79.9% ACN, 0.1% formic acid, 20% ultra-high purity water (MilliQ)] over 125 min and then to 80% solvent B over 5 min, at a flow rate of 300 nL min^-1^. Solvent A (v/v) consisted of 0.1% FA in high-purity water (MilliQ). Mass spectra were acquired in positive ion mode using a top-20 data-dependent acquisition (DDA) method on a Q-Exactive mass spectrometer (Thermo Fisher Scientific). A full MS scan was performed at 70,000 resolution (at m/z 200) with a scan range of 380–1800 m/z, followed by an MS/MS scan at 17,500 resolution (at m/z 200). For MS/MS fragmentation, higher energy collisional dissociation (HCD) was used with a normalized collision energy (NCE) of 27%. Dynamic exclusion was set to 20 seconds. Raw data were searched with the SEQUEST algorithm present in Proteome Discoverer (v1.3, Thermo Fisher Scientific). Proteins were annotated based on version 6.1 of the CC-4532 *C. reinhardtii* strain genome (Craig et al., 2022) retrieved from the Phytozome database (v13; Goodstein et al., 2012). Peptides were matched against this database plus decoys, considering a significant hit when the peptide confidence was high (FDR-adjusted *p*-value <0.01) and the Xcorr threshold was established at 1 per charge (2 for +2 ions 3 for +3 ions, etc.). The variable modifications were set to acetylation of the N-terminus and methionine oxidation, with a mass tolerance of 10 ppm for the parent ion and 0.8 Da for the fragment ion. The number of missed and nonspecific cleavages permitted was two. There were no fixed modifications, as dynamic modifications were used. The identified proteins were quantitated based on total ion count and normalized using the normalized spectral abundance factor (NSAF) strategy (Paoletti et al., 2006). All the MS/MS spectra of the identified proteins and their meta-information were further uploaded to the ProteomeXchange repository (project accession: PXD062167).

### ATP quantification

Cells, harvested as described above for metabolite profiling, were resuspended in 50 µL of ice-cold phosphate buffer (PBS, pH 7.5) and transferred to 2 mL reaction tubes. The samples were homogenized with a ball mill (30 Hz, 5 min) using two 3 mm glass beads to break the cell pellets and centrifuged at 20 000 *g* for 5 min at 4 °C. The ATP/ADP Ratio Assay Kit (MAK135; Sigma-Aldrich, St. Louis, MO, USA) was used for ATP quantification based on an external calibration curve. Bioluminescence was measured using a Plate reader (BioTek Synergy HTX; Agilent).

### Starch quantification

Total starch was quantified from the pellets collected during the GC-MS extraction using a commercial starch assay kit (MAK522, Sigma-Aldrich) according to the instructions of the manufacturer.

### Protein quantification and western blotting

Total soluble protein extracts were prepared from pellets collected during metabolite extraction. Proteins were extracted at 4 °C in 150 µL of lysis buffer containing 7 M urea, 2 M thiourea, 18 mM Tris-HCl, 14 mM Trizma base, 4% (w/v) CHAPS and 0.2% (v/v) Triton X-100, to which 20 µL of protease inhibitor cocktail (Complete Mini, Roche Diagnostics), dissolved in water as per the manufacturer’s instructions, were added. After 5 min at 4 °C, 3 µL of 1 M dithiothreitol (DTT) was added and the protein extracts were stirred for 30 min at 4 °C before centrifugation (20 000 *g*, 10 min, 4 °C). Protein concentrations were measured from the resulting supernatants according to (Bradford, 1976) using bovine serum albumin as standard.

Proteins were separated by 1D SDS-PAGE on 12 % poly-acrylamide gels and western blotting was conducted as previously described (Roach, 2020). Protein loading was adjusted based on the cultures OD_720_, resulting in an average amount of 4.8 ± 0.5 µg proteins loaded per well.

### Chlorophyll Fluorescence and P700 measurements

For transient near-infrared absorption of PSI reaction center chlorophyll (P700) and chlorophyll fluorescence measurements, 3 mL of culture was transferred to a cellulose acetate filter with 5 µM pore size (Sartorius, Göttingen, Germany) using a vacuum pump and Buchner funnel. The filter and cells were placed between a transparent plastic film for immediate measurements using the leaf clamps of a DUAL PAM (Walz, Effeltrich, Germany). Cells were first exposed for 3 min to LL and then 3 min to HL. At the end of each 3 min, an 800 ms saturating light pulse of 7044 µmol m^-2^ s^-1^ was given to calculate ETR of PSII and PSI, and NPQ, according to the equations provided in (Schreiber et al., 1986; Klughammer and Schreiber, 1994) prevent deactivation of the CBB, cells were not dark treated. Maximum fluorescence (F_m_) under LL was used for F_m_° in calculating the quenching that occurred under HL.

### Photoinhibition of PSII

Cells were harvested at an OD_720_ of 0.21 ± 0.01 by transferring 10 mL of the culture to 50 mL Erlenmeyer flasks. We used 2 mM lincomycin as an inhibitor of PS II repair, which was not added to the controls. After 30 min of dark acclimation, cells were re-incubated under their original light conditions for 30 min. Before the light treatment, 2.5 mM bicarbonate (NaHCO_3_) was added only to LL-FL- and HL-CO_2_ cells. A 2-mL aliquot of culture was used to measure the F_v_/F_m_ with an AquaPen-100 (PSI) after 30 min recovery in the dark.

### Statistical analysis

All statistical analyses were performed using R (R Core Team, 2024)(v4.4.1, R Core Team, 2024). For all univariate datasets, pairwise comparisons of the estimated marginal means were conducted using the ‘emmeans’ package, on log_2_ transformed data after fitting a linear model, and *p*-values were adjusted for multiple comparisons using the FDR correction (Benjamini and Hochberg, 1995). Differential protein and metabolite accumulation were assessed, after log_2_ transformation, using the Limma package (Ritchie et al., 2015) using a threshold set to 0.05 after global *p-*value adjustment for multiple comparisons using the FDR correction, unless otherwise specified. Sample size was at least 4 for all measurements. Heatmaps were generated using the ComplexHeatmap package (Gu et al., 2016), whereby hierarchical clustering was based on the Euclidean distance and Ward’s clustering method, and additional figures were prepared using the ggplot 2 package (Wickham, 2016).

## Acknowledgements

We thank Bettina Lehr, Christine Rossetti, Laurie Proes and Otto Dämon (University of Innsbruck) for excellent technical assistance. We thank Laurent Marquer for providing access to the fully motorized Olympus (BX53) microscope with the cellSens imaging software (ver.3.2) and deep-learning technology and Andreas Holzinger for insightful discussions regarding the cellular ultrastructure. Finally, we thank Suna Chiara Tarhan for the design of Figure 3.

## Author contributions

TR and AP planned and designed the research. TR, IK and WW contributed resources. SZ and LAS performed the proteomic analyzes with guidance from AG, PK and WW. AP, EAr and TR conducted all other experiments. EAr, AP and TR analyzed and interpreted the data. EAn, EG contributed new analytical and computational tools. AP, EAr and TR drafted the manuscript, which was revised and approved by all authors.

## Supplementary data

**Table S1.** Proteomics dataset

**Table S2.** Statistical evaluation of the proteomic data

**Table S3.** Proteins differentially accumulated depending on CO2 availability under the different light regimes.

**Table S4.** Metabolite profiling dataset

**Table S5.** Statistical evaluation of the metabolite profiling data

**Figure S1.** Influence of light intensity on gross oxygen production and light-enhanced dark respiration.

**Figure S2.** Influence of growth condition on maximum photosystem II efficiency, as measured with chlorophyll fluorescence (F_v_/F_m_).

**Figure S3.** Influence of growth condition on total photosynthetic electron flow, as measured with photosystem I (P700) reduction in the dark.

**Figure S4.** Influence of CO_2_ availability and light conditions on the proteome profiles, as visualised by principal component analysis.

**Figure S5:** CO_2_ availability differentially influenced protein accumulation depending on the light condition.

**Figure S6:** Light regimes differentially influenced protein accumulation.

**Figure S7.** Influence of growth conditions on rates of photoinhibition, as measured via decrease in maximum photosystem II efficiency (F_v_/F_m_).

**Figure S8.** Influence of growth conditions on the ratio of electron transfer rates of photosystem I to photosystem II (ETRI/ETRII) as a marker of cyclic electron flow.

**Figure S9.** Influence of growth conditions on the redox state of photosystem I reaction center chlorophyll (P700) in response to a saturating pulse.

**Figure S10.** Influence of growth conditions on metabolite profiles, as visualised by principal component analysis.

**Figure S11.** Influence of growth conditions on total fatty acids accumulation.

## Data availability

The data that supports the findings of this study are available in the supplementary material of this article and in the public ProteomeXchange repository (project PXD062167).

